# Modelling Actin-Microtubule Crosstalk in Migrating Cells

**DOI:** 10.1101/2025.04.09.647914

**Authors:** Pinaki Nayak, Anil Kumar Dasanna, Raja Paul, Heiko Rieger

## Abstract

Actin–Microtubule crosstalk regulates the polarity and morphology of migrating cells and encompasses mechanical interactions, mediated by crosslinkers, molecular motors, and cytoskeletal regulators. Recent experiments indicate that local microtubule depolymerization promotes local actomyosin retraction, whereas local microtubule polymerization promotes local actin-polymerization. Based on these observations, we develop a computational whole-cell model involving dynamic microtubules interacting mechanically and chemically with an active cell boundary. Specifically, the tips of microtubules send signals for local expansion or contraction to the active cell boundary, depending on whether they are in the growth or shrink phase. A rich, self-organized, dynamic behavior emerges, characterized by the repositioning of the microtubule-organizing center relative to the nucleus and the direction of migration. This also includes a variety of migration patterns, cell morphologies, and complex responses to obstacles in microfluidic and obstacle park environments. We demonstrate that microtubule length and cell boundary stiffness have a significant impact on these features, highlighting the need for new experimental investigations. Thus, the model provides a unified framework that explains a wide range of experimental observations and setups where actin-microtubule crosstalk plays a crucial role.

## INTRODUCTION

Cell migration is primarily driven by forces generated at the actin cortex underlying the cell membrane. The onset of migration requires the cell to be polarized by forming a protruding front edge and a contracting rear edge [1]. Protrusions at the front originate from the increased actin polymerization supported by the focal adhesions formed in contact with the extracellular matrix [2–4]. Membrane retraction at the rear is realized inside a cell by contractile forces arising from myosin activity and dissolution of focal adhesions [2, 5, 6]. The formation of protrusion or membrane retraction is guided through the reorganization of the cytoskeleton by the delivery of molecular regulatory signals. Microtubules (MT) are known to play an important role in the distribution of these regulatory signals leading to cell polarization during migration [7–9]. The tips of growing MTs reach the protruding front edge of the cell to deliver actin polymerization signals that stabilize the protrusions [6, 10, 11]. MT depolymerization induces the activation of RhoA, which increases myosin-II activity, increasing contractility and cell membrane retraction [12–14]. Differential stability of MTs at the front and rear edge thus leads to symmetry breaking and polarization of the cell [6, 15]. MTs have also been suggested to play a critical role in the modulation of cell shape and stabilization or retraction of protrusions when the cell is navigating through obstacles, thereby dictating the cell migration path [6]. Centrosomes, being the primary MT organizing centers (MTOC) in animal cells, can guide the choice of front and rear edge of the cell, via their preferential position with respect to the nucleus. The centrosomal position anterior or posterior to the nucleus has been investigated in various cells, proposing mechanisms that may guide this choice. Cells undergoing mesenchymal migration are characterized by the formation of nascent focal adhesions with the extracellular matrix at the leading edge and the rupture of aged focal adhesions at the rear edge [16–18]. The centrosome is placed ahead of the nucleus more often in cells migrating through stiff extracellular matrices [19]. Fast-moving amoeboid cells, characterized by reduced focal adhesions, are typically known to position their centrosome posterior to the nucleus during migration [20]. Interestingly, recent experimental studies suggest that leukocytes can alternate between a centrosome forward or nucleus forward configuration while navigating a congested microenvironment [21]. Cells may alter the position of the centrosome to modify the distribution of MTs, which in turn helps coordinate the polymerization and contraction signals in the actin cortex that drive cell movement. As MTs grow, they can extend toward the cell membrane or nucleus and undergo buckling. This buckling generates a pushing force that pushes the centrosome away from the point of contact with either the cell or nuclear membrane [22–24]. MTs can also slide along the cell membrane or nucleus [25]. Dynein motors present at the cortex or nuclear membrane attach to the sliding MTs and walk towards their minus-end, effectively pulling the MTs and the centrosome [22, 26–28]. The position of the centrosome within a moving cell results from a complex balance of forces, which are generated through the MTs’ interactions with the centrosome, along with membrane remodeling driven by polarity signals from the MTs to the cortex. This raises several questions: How does the centrosome position itself anterior or posterior to the nucleus during migration? How do changes in MT dynamics influence the centrosome’s position? And what impact do these alterations in the MT network have on actin-MT interactions and the cell’s overall migration?

Phenomenological and mechanistic acto-myosin models of cell migration have been the subject of extensive study [29–38]. Models that focus on the microscopic details of actin polymerization and myosin motor activity are computationally intensive and challenging to generalize for studying cell migration in both two (2D) and three dimensions (3D) [39, 40]. On the other hand, phenomenological models fail to capture the details of intracellular processes critical for symmetry breaking and force generation at the cortex [33, 41, 42].

While it is well established that MT-actin crosstalk plays a crucial role in cell migration and other cellular functions [43], a mechanistic whole-cell model integrating MT dynamics and actin-generated forces to explore the self-organization of centrosome positioning, cellular shape changes, and migratory behavior is still lacking. In this work, we therefore introduce a phenomenological model based on the experimental observations reported in [6, 7, 10, 13, 15], which correlate MT growth and shrinkage with actin polymerization, depolymerization, and contraction. We examine how the position of the centrosome and the direction of migration are influenced by MT dynamics. Additionally, we explore how cells might leverage MT-actin crosstalk to switch between persistent migration and diffusive movement. Finally, we investigate how the positioning of the centrosome, either anterior or posterior to the nucleus, may guide the cell’s path when navigating narrow channels.

## DESCRIPTION OF THE MODEL

A mechanistic model of cell migration primarily involves membrane protrusion and retraction, which are coupled to the actomyosin cortex and actin cytoskeleton. This process includes actin polymerization, myosin-driven membrane contraction, polarization of actin filaments, focal adhesion kinetics, and membrane surface tension. Research has shown that MT depolymerization regulates actomyosin contraction through the modulation of Rho GTPase signaling pathways [6]. On the other hand, MT polymerization can regulate actin polymerization and the expansion of protrusions by transporting intracellular cargo and signaling molecules to the leading edge of migrating cells [6, 10, 43]. We use these two observations as key components to develop a mechanistic whole-cell model of MT-actin crosstalk during cell migration.

We focus on two-dimensional mesenchymal cell migration, using the following basic model elements: 1) bead-spring loops to represent the semi-flexible boundary of the cell and its (circular) nucleus, 2) bead-spring semiflexible polymers for dynamic MTs (MTs), which can grow and shrink at their plus ends through the addition and removal of beads, respectively. These MTs are anchored at the MT organizing center (MTOC), which we assume to coincide with the cell’s centrosome, and 3) growing MTs exert pushing forces on the membrane when they come into contact, while dyneins, anchored at the cell and nucleus membranes, can attach to MTs and generate pulling forces on them. A sketch of these standard parts of our model (c.f. [44]) is shown in Fig. 1 and all mathematical details are given in the appendix. As new model elements, we incorporate MT-actin crosstalk by introducing a regulatory cue that modulates increased actin polymerization at the cortex or membrane contraction due to myosin-II activity, which is locally delivered to the cortex at the MT tips. Shrinking MTs deliver cues for myosin activity, leading to local contraction, while growing MTs deliver cues for actin polymerization, resulting in local expansion. Instead of modeling the actin network explicitly, we represent local contraction and expansion through their effective action on the individual beads that represent the cell boundary. First, actin-MT crosstalk due to shrinking MTs leads to myosin-generated stress in the actin cortex, causing membrane retraction via the RhoA GEF LFc signaling pathway [6]. We model this event chain by introducing an effective inward force on a membrane bead that is close to the tip of a shrinking MT. The net inward force on a membrane bead is proportional to the number of shrinking MT tips in the vicinity of the membrane bead (see Fig. 1 and the appendix for the mathematical details). Second, actin-MT crosstalk due to growing MTs leads to the formation of membrane protrusions or lamellipodia due to actin polymerization against a membrane. This induces a retrograde actin flow toward the nucleus, which is opposed by focal adhesions connecting actin filaments across the membrane to the substrate. To describe this sequence of events, we examine a model of lamellipodial protrusion guiding confined cell migration [33]. In this model, the local actin polymerization force acting on a segment of the cell membrane is linked to both the actin polymerization rate and the average orientation of actin filaments near the membrane. [33, 41, 45, 46]. Consequently, the actin polymerization force generates an outward-directed velocity component on the membrane beads, which is influenced by the polymerization signals transmitted from the growing microtubule (MT) tips to the actin cortex (c.f. Fig. 1). To incorporate actin-MT crosstalk, we assume that the outward velocity imparted to the membrane bead along the outward normal is proportional to the number of growing MT tips near the membrane bead (see the appendix for the mathematical details). Additionally, myosin-generated contraction forces, mediated by actin filaments between the protrusion’s leading edge and the nucleus, counteract the expansion of the protrusion, resulting in an effective elastic coupling between the position of the nucleus and the protrusion edge [33]. We model this elastic coupling by linking the beads of the nucleus boundary to those of the cell boundary with elastic springs, as illustrated in Fig. 1 (see the appendix for the mathematical details).

**FIG. 1.**
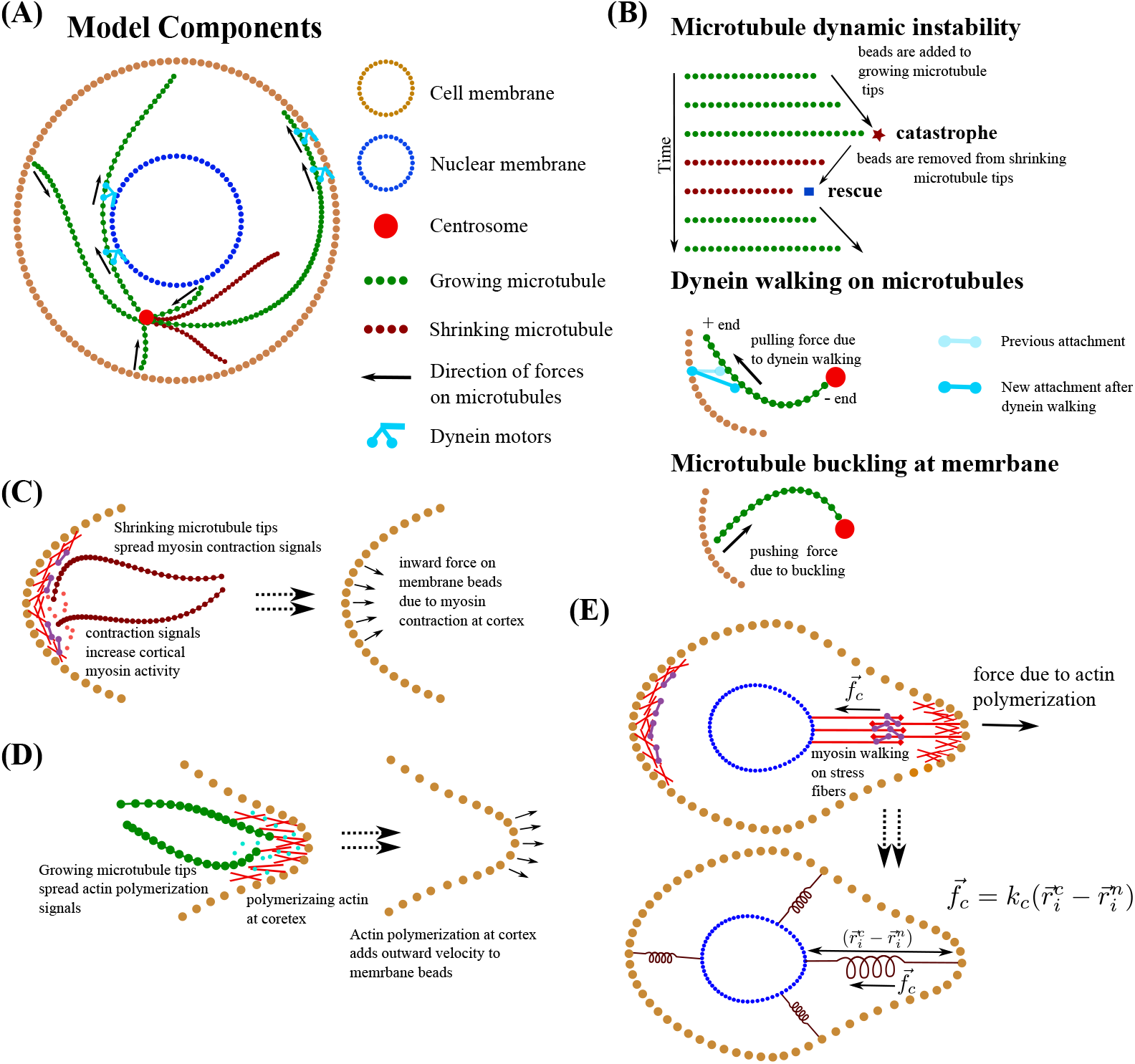
Schematic of the model: **(A)** A sketch of the migrating cell, consisting of membrane, nucleus, centrosome, MTs, actin and dynein motors. **(B)** A MT in its growth phase adds beads to its plus-end tips, while shrinking MTs have beads removed from the plus-end tips. A growing MT can undergo catastrophe and begin shrinking, while a shrinking MT can experience rescue and start growing again. Dynein, anchored at the cell and nuclear membranes, attaches to the MTs and moves toward their minus ends, applying pulling forces on the centrosome. The steric interactions between MT beads and the membrane cause MT buckling, resulting in pushing forces on the centrosome. **(C)** Shrinking MTs transmit myosin contraction signals to the actin cortex, where myosin activity generates an inward-directed contraction force on the membrane beads. **(D)** Growing MT tips transmit actin polymerization signals to the cell cortex, where actin polymerization drives an outward velocity of the membrane beads. **(E)** Myosin contractility on the overlapping actin filaments of the cell and nuclear membranes creates a linear elastic coupling between the nuclear and cell membrane beads, which counteracts the protrusion forces on the cell membrane.

## RESULTS

### Migrating cell centrosome can lead or trail nucleus depending upon MT length

First, we examine how centrosome positioning and migration characteristics in our model are influenced by MT properties, such as the average MT length. The trajectories of the cell centroid indicate that for average MT lengths of *l*_*mt*_ = 9.3 − 16*µm*, the cell exhibited directed migration, whereas for an average MT length of *l*_*mt*_ = 3*µm* the cellular trajectories showed random motion (Fig. 2**A**; see Supplementary Movie **1-2** [47]).

**FIG. 2.**
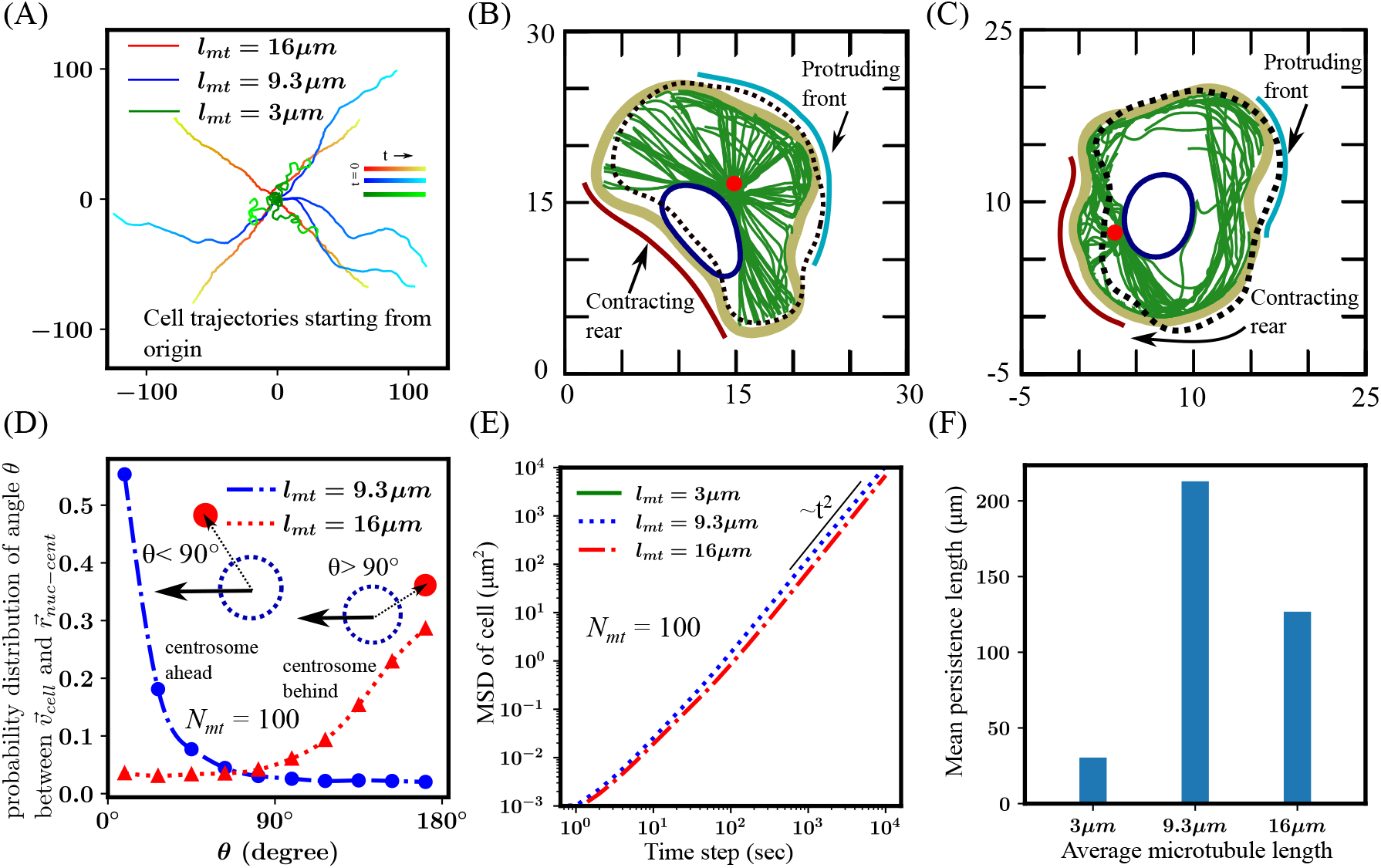
Centrosome position depends on the MT length. **(A)** Cell trajectories corresponding to different average MT lengths show directed migration for average MT lengths *l*_*mt*_ = 3*µm*, 9.3*µm* (2*/*3 of cell diameter) and 16*µm* (greater than cell diameter). In contrast, for a shorter average MT length of *l*_*mt*_ = 3*µm*, the trajectories indicate random motion. **(B)** Simulation snapshots showing centrosome anterior to nucleus for *l*_*mt*_ = 9.3*µm*. Growing MT tips at the front deliver actin polymerization signals, driving protrusion formation. The cell rear retracts through an elastic coupling between the nucleus and the membrane. **(C)** Simulation snapshots show the centrosome positioned posterior to the nucleus (*l*_*mt*_ = 16*µm*). Growing MT tips accumulate at the cell front, where they provide actin polymerization signals to drive protrusion formation. Shrinking MT tips transmit myosin contraction signals along the cell rear, initiating actin cortex contraction and membrane retraction. **(D)** Probability distribution of the angle between the cellular direction of motion and the vector from the nucleus to the centrosome; *θ <* 90° indicates the centrosome is ahead of the nucleus, while *θ >* 90° indicates the centrosome is behind the nucleus. Regular MTs (*l*_*mt*_ = 9.3*µm*) position the centrosome ahead of the nucleus, while longer MTs (*l*_*mt*_ = 16*µm*) position the centrosome behind the nucleus. **(E)** Mean square displacement of cells exhibits a *t*^2^ dependence, indicating ballistic migration for *l*_*mt*_ = 9.3 − 16*µm*. **(F)** Mean persistence length of migrating cell for different average MT lengths. Results indicate that cell migration is highly persistent for *l*_*mt*_ = 9.3 − 16*µm*.

We then focused on the MT lengths that resulted in the directed migration of the cell. Our results showed that for an average MT length of *l*_*mt*_ = 9.3*µm* (which is 2/3 of the cell diameter, henceforth referred to as “regular MT”), the centrosome was predominantly positioned ahead of the nucleus (Fig. 2 **B,D** (blue line)). When the average MT length was increased to *l*_*mt*_ = 16*µm* (greater than the cell diameter, henceforth referred to as “long MT”), the centrosome preferentially remained behind the nucleus in the direction of migration (Fig. 2**C,D** (red line)). To understand this preferential positioning of the centrosome in both cases, we examined the MT dynamics. MTs radiated from the centrosome in all directions, extending to both the cell and nuclear membranes. However, the presence of the nucleus obstructed the MTs from reaching the portion of the cell membrane behind it. Regular MTs were not long enough to slide along the nuclear membrane and extend to the rear portion of the cell membrane (Fig. 2**B**). The growing MTs that reach the cell membrane have most of their length within the cytoplasm, with only a small portion near the MT tips remaining in close contact with the cell membrane (Fig. 2**B**). The MT tips deliver actin polymerization signals to the membrane region near their tips. When an MT undergoes catastrophe, its tip recedes from the membrane. As a result, shrinking MTs do not remain in contact with the membrane long enough to spread contraction signals continuously. Consequently, actin polymerization signals are only spread in regions of the membrane with a high density of MT tips, promoting the formation of protrusions. Areas of the membrane lacking MT tips are predominantly influenced by the nucleus-to-cell membrane elastic coupling, forming the contractile rear end of the cell. This asymmetry in the distribution of actin polymerization signals breaks the symmetry of the stationary cell. The regions receiving signals from growing MTs form the protruding front end, while areas devoid of MT tips form the rear. As a result, the cell migrates with the centrosome near the cell center and the nucleus positioned behind it towards the rear end of the cell. For regular MTs, we find that the probability distribution of the angle between the cell’s direction of motion and the vector from the nucleus to the centrosome peaks near zero (Fig. 2**D** (blue line)), corroborating the result that the centrosome remains positioned ahead of the nucleus throughout migration.

Cells with long MTs (*l*_*mt*_ = 16*µm*) are polarized with the centrosome positioned posterior to the nucleus. MT tips extending from the centrosome reach either the cell membrane or nuclear membrane and can glide along these membranes to eventually reach the distal end of the cell, behind the nucleus (Fig. 2**C**). Since the average MT length exceeds the cell’s diameter, most growing MTs glide along the cell or nuclear membrane until they reach the distal end of the cell (Fig. 2**C**). Eventually, the tips of these growing MTs accumulate at the distal end of the cell membrane, where they promote actin polymerization signals. This leads to a higher concentration of actin polymerization cues at the membrane region farthest from the centrosome. In contrast, long MTs that reach the distal cell membrane may undergo catastrophe. These MTs often glide along or remain close to the cell membrane (Fig. 2**C**). When catastrophe occurs, the MTs shrink toward the centrosome, spreading contraction signals along the cortical region they pass through. As the MT tips recede, they move away from the distal cell mem-brane and get closer to the centrosome. Consequently, the contraction signals from the shrinking MT tips are primarily concentrated near the centrosome. This process polarizes the cell, with protrusions forming at the distal membrane (farthest from the centrosome) and retraction occurring at the proximal membrane (closer to the centrosome) due to elevated cortical myosin activity. As a result, the cell migrates with the centrosome positioned posterior to the nucleus. For long MTs, the probability distribution of the angle between the cell’s direction of motion and the vector from the nucleus to the centrosome peaks near 180° (Fig. 2**D** (red line)), suggesting that the centrosome largely remains posterior to the nucleus during cell migration [5, 20].

Next, we characterize the persistence of migrating cells for various average MT lengths, which correspond to different centrosome positioning. The mean square displacement (MSD) of the cell centroid, scaled by the square of time, suggests ballistic motion (*MSD* ∝ *t*^2^) (Fig.2**E**). This indicates that the cell can move in a ballistic mode regardless of whether the centrosome is positioned anterior or posterior to the nucleus, as long as the cell remains polarized. We also examined how the persistence length of migrating cells changes with different MT lengths. For short MTs (*l*_*mt*_ = 3*µm*), the mean persistence length was small (≈ 25*µm*) indicating that the cell frequently deviated from its direction of migration over short distances (Fig. 2**F**). In contrast, the mean persistence length was significantly higher for regular and longer MTs (*l*_*mt*_ = 9.3 and 16*µm*, respectively), suggesting more persistent migration. Interestingly, the persistence length was greatest for the regular MT length of *l*_*mt*_ = 9.3*µm*, indicating that cells with this MT length exhibit the most consistent directionality during migration.

### Variation of MT numbers affects cell polarization and migration persistence

Next, we investigate how variations in MT number influence a cell’s migration characteristics. Experimental studies suggest that changes in MT abundance can either enhance or impede directed locomotion [48–50]. For instance, increased MT numbers have been linked to greater persistence in dendritic cells and cancer metastasis [48, 50]. We ran simulations with different numbers of regular and long MTs, initially focusing on regular MTs (*l*_*mt*_ = 9.3*µm*), with MT counts ranging from *N*_*mt*_ = 25 to 200. As MT numbers increase, the centrosome becomes more prominently positioned ahead of the nucleus, in the direction of migration (Fig. 3**A**). With more MTs nucleating from the centrosome, a greater number of MTs reach the cell front, enhancing the delivery of actin polymerization signals. Consequently, cells with more MTs exhibit increased actin polymerization activity at the front. However, due to the relatively short length of regular MTs, most are unable to circumvent the nuclear membrane and extend to the membrane region behind the nucleus. This limits the actin polymerization signals in the rear cell membrane. Regular MTs only have a small portion of their length near the tip in proximity to the cell membrane (Fig. 2**B**), and when they undergo catastrophe, their tips retract from the membrane. Consequently, acto-myosin contraction signals are not efficiently transmitted to the actin cortex beneath the membrane. Our findings show that as MT numbers increase, the angle between the cell’s direction of motion and the vector from the nucleus to the centrosome tends to be smaller (i.e., the centrosome is more often ahead of the nucleus) (Fig. 3**A**). This suggests that higher MT numbers contribute to more stable protrusions at the membrane regions near the centrosome. The rear of the cell, receiving fewer polymerization signals, contracts through elastic coupling between the nucleus and the membrane. As a result, the cell tends to break symmetry and migrate, with the centrosome positioned ahead of the nucleus.

**FIG. 3.**
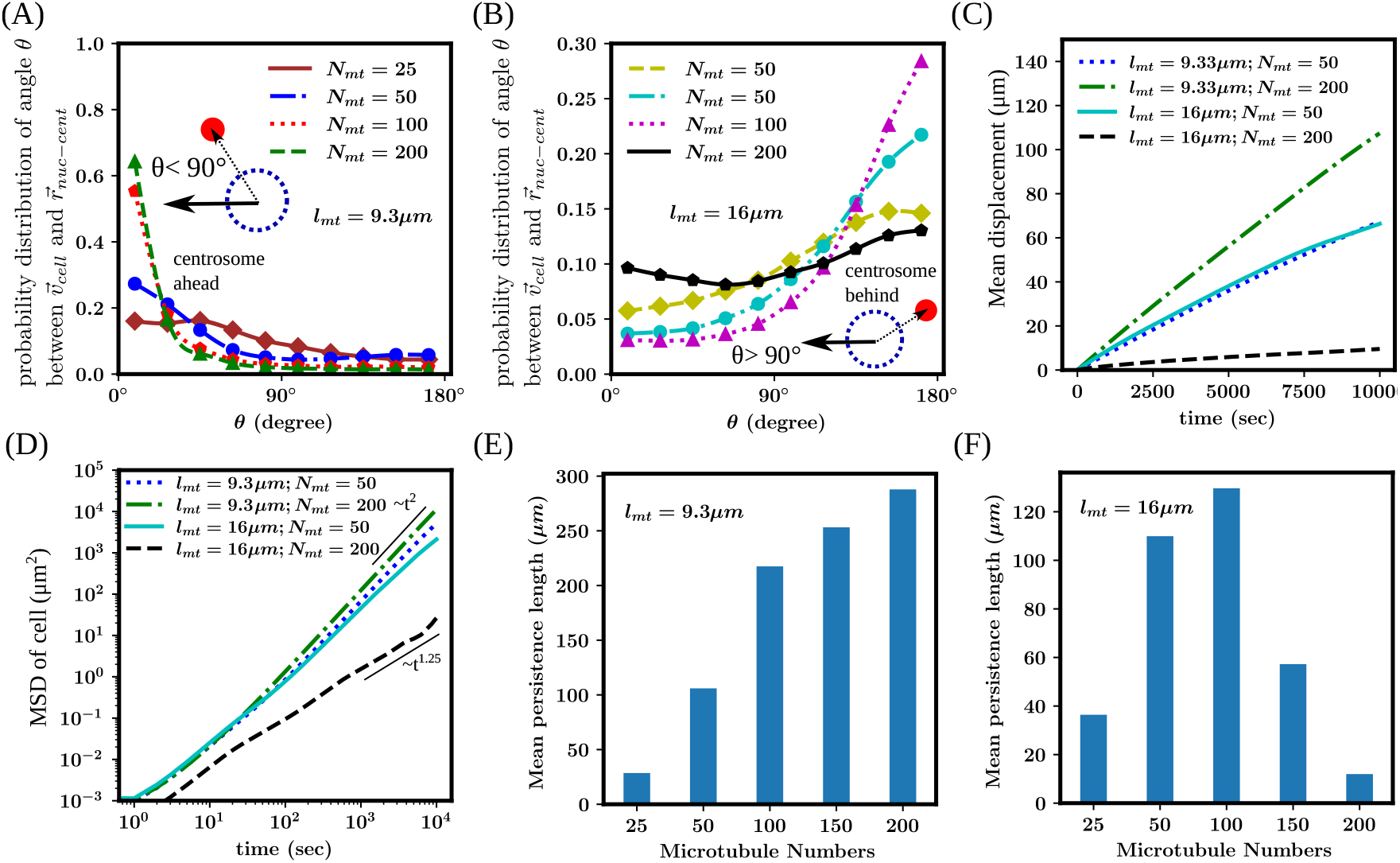
MT number affects persistence of cell migration. **(A-B)** Probability distribution of the angle between the direction of motion and the nucleus to centrosome vector for *l*_*mt*_ = 9.3*µm* and 16*µm*, at different MT numbers. **(C)** Mean displacement of the cell centroid as a function of time for varying MT numbers and average lengths. **(D)** Mean square displacement of the cell over time for varying MT numbers and average lengths. An increase in the number of MTs, with an average MT length of *l*_*mt*_ = 16*µm*, causes the cell to transition from ballistic motion (*MSD* ∝ *t*^2^) to super diffusive motion (*MSD* ∝ *t*^1.25^). **(E)** Persistence length of migration for various MT numbers with *l*_*mt*_ = 9.3*µm*. persistence length increases steadily as the MT number increases. **(F)** Persistence length of migration for various MT numbers with *l*_*mt*_ = 16*µm*. Persistence length of the cell initially increases with MT number but decreases when the MT number becomes high.

Our results demonstrate that cells with regular MTs exhibit more robust migration as the MT number increases. The displacement of the cell centroid was consistently greater for cells with *N*_*mt*_ = 200 compared to *N*_*mt*_ = 50 at all time points (Fig. 3**C**). Analysis of the mean squared displacement (MSD) of the cell centroid revealed ballistic movement (*msd* ≈ *t*^2^) for cells with *N*_*mt*_ = 50 − 200 (Fig. 3**D**). Furthermore, the persistence length of migrating cells increased with MT numbers (Fig. 3**E**), indicating that cells with more MTs maintained their polarization and were less likely to change the direction of migration. [48].

Next, we investigate the effect of MT numbers for long MTs (*l*_*mt*_ = 16*µm*). For low to intermediate MT numbers (*N*_*mt*_ = 25 − 100), the growing MT tips eventually reach the distal end of the membrane, where they deliver actin polymerization signals and shrinking microtubules spread contraction signals near the rear end of the cell. As the number of MTs increases, the probability distribution of the angle between the migration direction and the nucleus-to-centrosome vector shifts toward 180° (Fig. 3**C**), indicating a stronger cell polarization. However, when the MT number becomes very high, polarity is reduced. High MT numbers result in more growing MT tips near the rear of the cell membrane, which weakens the polarity gradient. Consequently, both ends of the membrane receive significant actin polymerization signals. For *N*_*mt*_ = 200, the cell no longer maintains a distinct anterior or posterior centrosome configuration during migration. The probability distribution of the angle between the migration direction and the nucleus-to-centrosome vector becomes more even, with no clear peak at any angle (Fig. 3**C**(black line)). With a very large number of MTs, the cell continuously shifts its front and rear based on the actin polymerization signals received along each segment of the membrane, whether proximal or distal to the centrosome.

Analysis of the cell trajectories reveals that cells with long MTs exhibit more robust movement when the MT numbers are between *N*_*mt*_ = 50 − 100. The mean displacement of the cell centroid shows that cells with *N*_*mt*_ = 50 travel greater distances over time compared to those with *N*_*mt*_ = 200 (Fig. 3**C**). The mean squared displacement of the cell centroid suggests that cells with *N*_*mt*_ = 50 migrate ballistically (*msd*≈ *t*^2^), while cells with *N*_*mt*_ = 200 exhibit super-diffusive motion (*msd* ≈ *t*^1.25^) (Fig. 3**D**). The persistence length of migrating cells increases with MT numbers within the range of *N*_*mt*_ = 25 − 100. However, for *N*_*mt*_ = 200, the persistence length sharply decreases to a very low value (Fig. 3**F**). This suggests that the persistence of directional locomotion increases with MT numbers for cells with long MTs (*l*_*mt*_ = 16*µm*), consistent with findings from various experiments [48]. However, a very large number of MTs can impair cell locomotion when the centrosome is located behind the nucleus.

### Anterior centrosome position improves directed migration in obstacle parks

We further examine the relationship between centrosome positioning and cell migration in obstacle parks, as explored experimentally in studies, e.g., [51, 52]. These studies highlight that cells exhibiting directed migration on flat surfaces can become trapped when placed within obstacle parks. To understand how actin-MT crosstalk enables cells to navigate through restrictive geometries, we performed simulations with the cell placed in obstacle parks of varying obstacle sizes and spacings. To assess whether cell migration was directed or random, we calculated the local mean squared displacement of the cell centroid (Δ*R*^2^(*t*_*i*_)) at regular time intervals of 300 seconds and examined its scaling exponent *α* as a function of the time lag 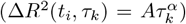 [51]. Additionally, we assessed the standard deviation of the velocity angle (Δ*ϕ*_*i*_) at the corresponding time intervals. Directed migration was defined as *α >* 1.7 and Δ*ϕ*_*i*_ *<* 0.9 (see appendix for details) [51].

We first analyzed the trajectories of freely migrating cells. Our results indicate that freely migrating cells undergo significant directed migration phases, both for regular and long MTs (Fig. 4**A-B**). When the cells were placed in an obstacle park (obstacle radius *R*_*obs*_ = 4*µm* with spacing Δ*d* = 20*µm*), the frequency of directed migration decreased (Fig. 4**C-D**; see supplementary movie **3**[47]). However, cells with regular MTs exhibit more directed migration phases in the obstacle park compared to those with long MTs. In cells with regular MTs, the centrosome is positioned ahead of the nucleus, allowing forward-growing MTs to explore alternative paths when encountering obstacles. This enables the cells to extend protrusions into gaps between obstacles and navigate through without fully changing direction. In contrast, cells with long MTs position their nucleus ahead of the centrosome, which can block MT extension toward adjacent pores when an obstacle is encountered. As a result, cells with long MTs tend to avoid obstacles and narrow pores, leading to a significant decrease in directed migration phases. Increasing the obstacle size further (*R*_*obs*_ = 5*µm*) reduces the frequency of directed migration for both regular and long MT cells. However, cells with regular MTs are still able to maintain more directed migration compared to long MT cells, which are more likely to become trapped (Fig. 4**E-F**; see Supplementary movie **4**[47]). This indicates that when there is enough space for cells to pass through, they can maintain directed migration. Moreover, the mean displacement of the cell centroids revealed that cells with regular MTs, where the centrosome is positioned anterior to the nucleus, traveled greater distances in the obstacle park, regardless of obstacle size (Fig. 4**H**).

**FIG. 4.**
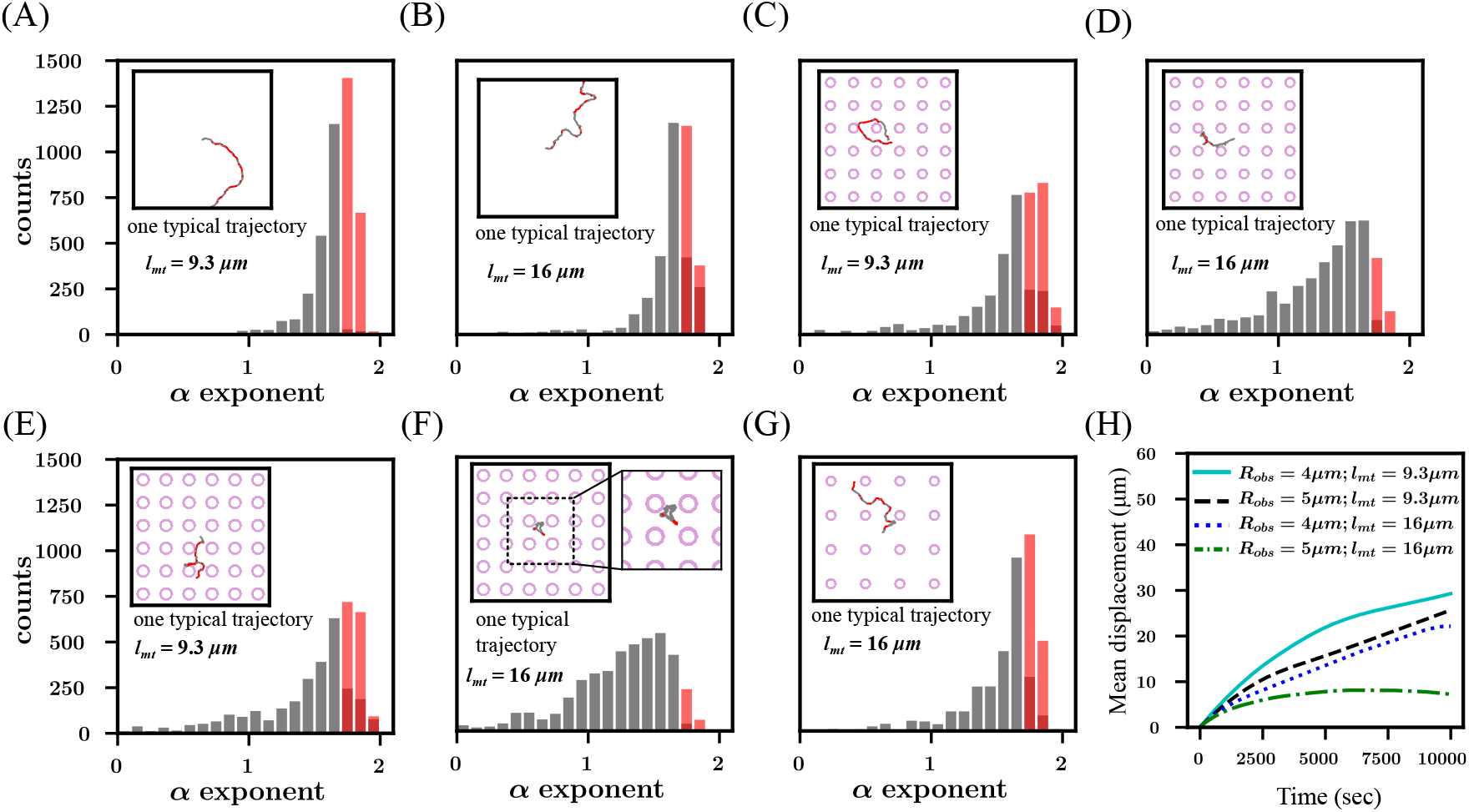
Cell migration through obstacles. **(A-B)** Distribution of local MSD exponent *α* for free cell migration for *l*_*mt*_ = 9.3*µm* and 16*µm* with 50 MTs. Grey bars represent counts for random migration, while red bars indicate counts for directed migration. **(C-D)** Distribution of *α* for cell migration in obstacle park with obstacle radius *R*_*obs*_ = 4*µm* and obstacle spacing Δ*d* = 20*µm* for *l*_*mt*_ = 9.3*µm* and 16*µm*. **(E-F)** Distribution of *α* with larger obstacles having radius *R*_*obs*_ = 5*µm* and obstacle spacing Δ*d* = 20*µm* for *l*_*mt*_ = 16*µm*. **(G)** Distribution of *α* with obstacle radius *R*_*obs*_ = 4*µm* and larger obstacle spacing Δ*d* = 30*µm* for *l*_*mt*_ = 9.3*µm* and 16*µm*. All trajectories shown in insets start at the center of the frame of size 60*µm ×* 60*µm*. Y axis range is same for Fig. A-G. **(H)** Mean displacement of the cell centroid over time for various obstacle sizes and MT lengths.

### Cell morphology altered by length and number of MTs

Next, we examined how the actin-MT crosstalk influences the morphology of the migrating cells, quantified by the cell’s aspect ratio and spread area. When MTs are long (*l*_*mt*_ = 16*µm*), the aspect ratio remains close to 1 for all MT numbers, suggesting the cell shape is nearly circular (Fig. 5**A,E**). In this configuration, the centrosome is preferentially positioned behind the nucleus, near the rear of the cell. MT tips extend along both the nuclear and cell membranes, reaching the opposite end of the cell, where they deliver actin polymerization signals that drive membrane protrusions. The region closer to the centrosome has a higher density of MTs, helping to resist membrane contraction. This balance leads to a roughly circular cell shape. However, when the average MT length is shorter (e.g. regular MT length with *l*_*mt*_ = 9.3*µm*), the aspect ratio varies between 1.5 and 3.5 for *N*_*mt*_ = 50 (Fig. 5**A,F**). As MT numbers increase (*N*_*mt*_ = 100, 200), the aspect ratio stabilizes around 1.5. This indicates that the cell shape deviates from a circular form when *l*_*mt*_ = 9.3*µm*. In this scenario, the centrosome remains positioned ahead of the nucleus, closer to the front of the cell membrane. However, the MTs fail to navigate around the nuclear membrane to reach the opposite end of the cell. Consequently, actin polymerization signals are not delivered to the membrane region behind the nucleus. Thus, membrane protrusions only occur at the front and side regions of the cell membrane, where MTs are able to grow. In contrast, the rear of the membrane receives fewer MTs and experiences contraction forces from the membrane-nucleus coupling. The combination of extension at the front and sides, alongside contraction at the rear, leads to the cell deviating from its circular shape.

**FIG. 5.**
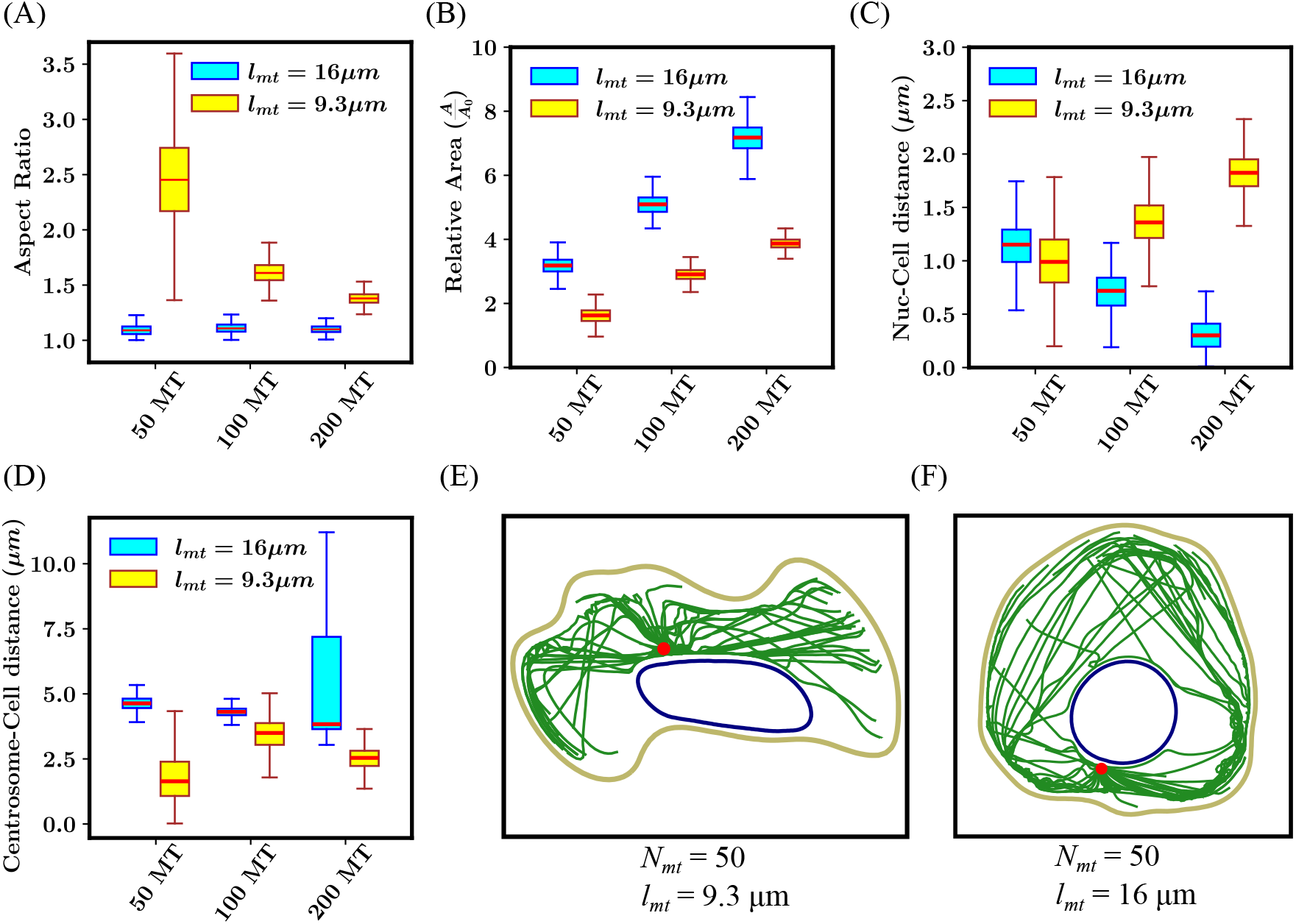
MT length and numbers influence cell morphology. **(A)** Aspect ratio of cells with various MT lengths and numbers. Cells with long MTs have an aspect ratio ∼ 1 (circular shape). Regular MTs having *l*_*mt*_ = 9.3*µm* lead to a high aspect ratio when the MT number is low (*N*_*mt*_ = 50). **(B)** Area of the migrating cell relative to initial area 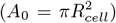 for various MT lengths and numbers. Cell area increases monotonically with MT numbers and average length. **(C)** Distance between the nucleus and the cell center for varying MT lengths and numbers. **(D)** Distance of centrosome from cell center for various MT lengths and numbers. **(E-F)** Snapshots of migrating cells for *l*_*mt*_ = 9.3*µm* and 16*µm* with *N*_*mt*_ = 50.

Next, we investigate the spread area of the cells for various MT numbers and lengths. Actin polymerization activity at the cortex driven by MT delivered polymerization signals increases the spread area of the cell. Our results indicated that cells with long MTs have a larger spread area as compared with cells with regular MTs (Fig. 5**B**). Regular MTs deliver actin polymerization signals only at the membrane regions close to the centrosome. However, long MTs deliver actin polymerization signals throughout the cell membrane, resulting in longer protrusions and increased spread area of the cell. As MT numbers increase, the actin polymerization signals at the actin cortex also increase. Thus, an increase in MT numbers corresponds to an increase in cell area for regular and long MTs.

Next, we analyze the position of the nucleus relative to the cell center. For cells with regular MTs, the nucleus remains closest to the cell center when *N*_*mt*_ = 50 (Fig. 5**C**). As the number of MTs increases, the distance between the nucleus and the cell center also increases. In these cells, the centrosome is positioned anterior to the nucleus. As the MT count rises, the pushing force exerted on the nucleus grows stronger, shifting the nucleus toward the rear of the cell and away from the center (Fig. 5**E**). In contrast, for cells with long MTs, the nucleus moves closer to the cell center as the number of MTs increases (Fig. 5**C**). In these cases, the centrosome remains positioned posterior to the nucleus. With more MTs, the increased pushing force on the nucleus results in its movement toward the cell center.

Finally, we examine the distance of the centrosome from the cell center. For cells with regular MTs, the centrosome remains closer to the cell center (Fig. 5**D**). When the centrosome is positioned anterior to the nucleus, our results indicate that it tends to stay near the cell center. The nucleus is slightly displaced toward the rear of the cell due to the pushing forces exerted by the MTs (Fig. 5**E**). Many regular MTs buckle at the cell membrane, and the resulting forces push the centrosome toward the cell center. For cells with long MTs, most of the MTs slide along the cell or nucleus membrane to the cell front. Consequently, the resultant pushing force on the centrosome is insufficient to keep it near the center. As a result, the centrosome remains toward the rear of the cell, while the nucleus moves closer to the center (Fig. 5**D,F**). As the number of long MTs increases, the pushing forces on the centrosome intensify, causing it to shift slightly closer to the center at *N*_*mt*_ = 100 as compared to *N*_*mt*_ = 50. However, with further increases in MT numbers, the cell loses its polarity, and the centrosome’s position becomes more random within the cell.

### MT actomyosin crosstalk influences cell path at Y junction

Inspired by experimental observations linking cell migration paths to centrosome positioning at Y-junctions [21], we placed our model cell in a comparable setup, as depicted in Fig. 6**A,B**. To replicate the chemotactic gradient applied in the experiments to drive forward migration, we impose a small forward velocity on the membrane beads. Initially, the channel widths are kept equal, larger than the nucleus diameter (*R*_*nuc*_ = 3*µm*), but smaller than the cell diameter (*R*_*cell*_ = 7*µm*), set at 10*µm*. We consider that a cell has moved into a channel when all of the cell has travelled at least 5.0 *µm* into a channel. Our results show that migrating cells with long MTs and the centrosome positioned behind the nucleus take significantly longer to enter the channel compared to cells with regular MTs and the centrosome ahead of the nucleus (Fig. 6**C**). For these equal-width channels, the probability of moving into either edge of the Y-junction is close to 50% for both regular and long MTs (Fig. 6**D,E**) and for regular MTs the cell took always the path that the centrosome entered first, in agreement with the experimental observations [21].

**FIG. 6.**
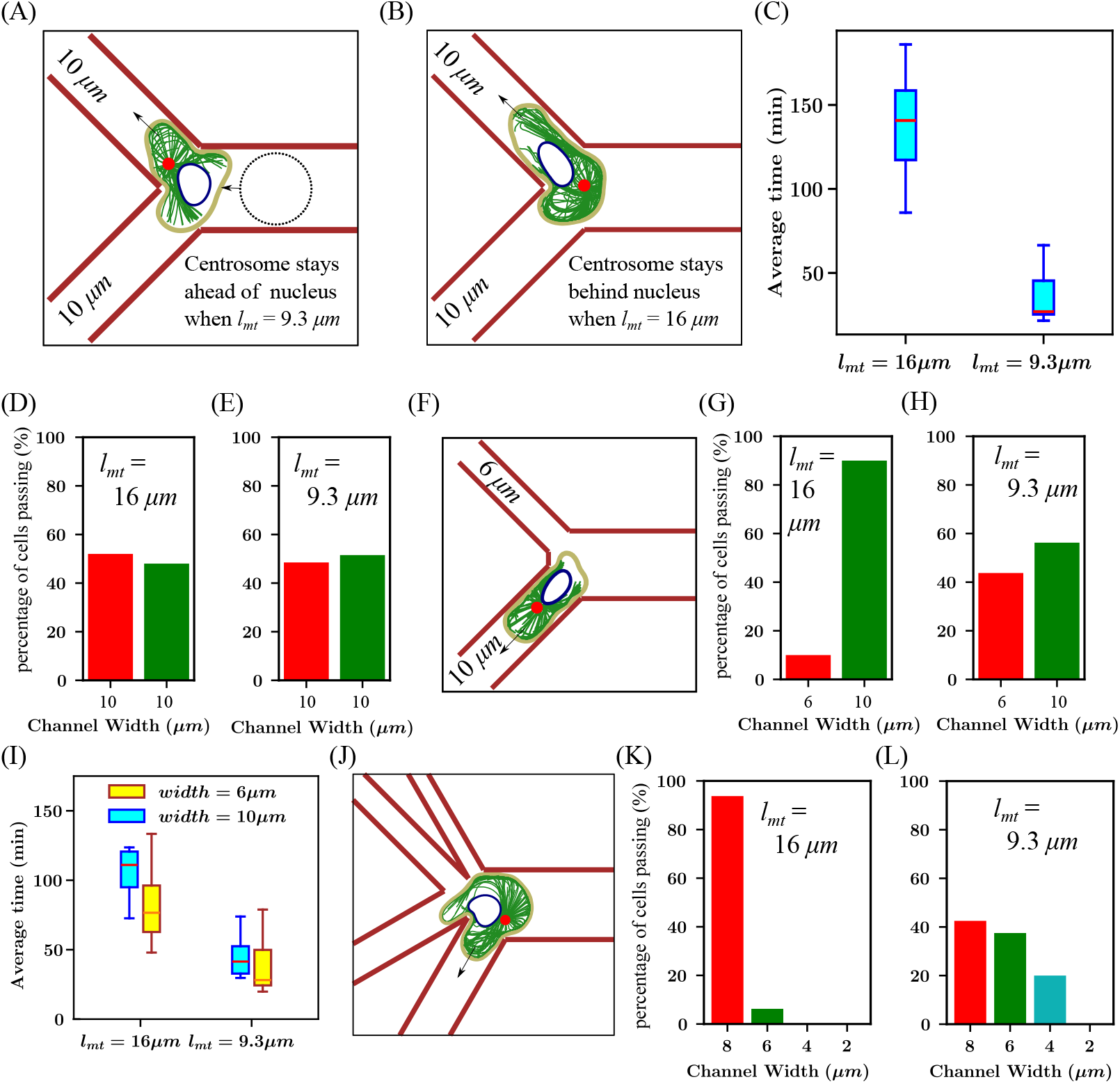
Cell migration in a Y-shaped channel. **(A-B)** Snapshots of cells navigating channels of equal widths (10*µm*) at the Y junction for *l*_*mt*_ = 9.3*µm* and 16*µm*. Dotted line in **A** shows initial position of cell and arrows indicate direction of migration. **(C)** Time taken by cells with *N*_*mt*_ = 100 to pass through Y junction with equal channel width 10*µm*. **(D)** Percentage of cells with long MTs (*l*_*mt*_ = 16*µm*), passing through each channel of width 10*µm*. **(E)** Percentage of cells with regular MTs (*l*_*mt*_ = 9.3*µm*), passing through each channel of width 10*µm*. **(F)** Snapshot of a cell at the Y junction with a wide pore (10*µm*) and a narrow pore (6*µm*).**(G)** Percentage of cells with long MTs (16*µm*), passing through wide pore (10*µm*) and narrow pore (6*µm*). **(H)** Percentage of cells with regular MTs (9.3*µm*), passing choosing wide pore (10*µm*) or narrow pore (6*µm*). **(I)** Time taken by cells with different MT lengths to move into a wide pore (10*µm*) or narrow pore (6*µm*). **(J)** Snapshot of a cell in a path choice device with four channels of varying widths.**K** Percentage of cells with long MTs (*l*_*mt*_ = 16*µm*), passing through each pore of different width in path choice device. **L** Percentage of cells with regular MTs (*l*_*mt*_ = 9.3*µm*), passing through each pore of different width in path choice device.

Next, we simulated cells facing a Y-junction with wide (10*µm*) and narrow (6*µm*) channels (Fig. 6**F**). For migrating cells with long MTs (*l*_*mt*_ = 16*µm*) and the nucleus ahead of the centrosome, cells prefer to move into the wider channel (Fig. 6**G**). This behavior is consistent with experimental observations suggesting that cells use their nucleus to gauge pore sizes and choose the path of least resistance [21]. Cells with regular MTs (*l*_*mt*_ = 9.3*µm*) and the centrosome ahead of the nucleus show a higher likelihood of moving into the narrow pore compared to cells with long MTs (Fig. 6**H**). However, these cells still have a higher probability of moving into the wider pore over the narrower one (Fig. 6**H**). This may be attributed to the fact that, with the centrosome ahead of the nucleus, growing MTs can easily extend into the protrusion within the smaller channel, facilitating actin polymerization. In contrast, cells with long MTs, where the nucleus is positioned ahead of the centrosome, experience blockage from the nucleus, preventing the MTs from reaching the smaller channel. As a result, membrane protrusions in the smaller channel are less stabilized and eventually retract.

Our results indicate that cells with long MTs take more time to move into either the narrow or wide channel compared to cells with regular MTs. However, when the width of the two channels is unequal, the time taken by cells with long MTs to choose a channel decreases (compare Fig. 6**C** and **I**). This suggests that when one path is much more restrictive than the other, cells with their nucleus ahead of the centrosome retract their protrusion from the restrictive path faster. When both the path choices are similar, the cell takes more time to retract its protrusion from one path and move completely into the other channel.

Finally, we examined the behavior of cells at a junction with multiple path choices. We simulated cells with varying MT lengths within a geometry containing four paths, each with different channel widths (Fig. 6**J**). Our results indicate that cells with long MTs (*l*_*mt*_ = 16*µm*) explore all available paths and only enter channels with wider pores (channel width = 8*µm* and 6*µm*), while no cells moved into channels with very narrow pores (channel width = 4*µm* and 2*µm*) (Fig. 6**K**; see Supplementary movie **5**[47]). The majority of the cells (≈ 90%) moved into the widest pore (width = 8*µm*), indicating a clear preference for the path of least resistance. In contrast, cells with regular MTs (*l*_*mt*_ = 9.3*µm*) did not show a strong preference for the widest pore, with about ≈ 40% of cells moving into the 6*µm* channel (Fig. 6**L**). Nearly 20% of cells moved into the 4*µm* channel, while no cells moved into the 2*µm* channel (see Supplementary movie **5**[47]). These results suggest that with the centrosome positioned ahead of the nucleus, MTs can extend further into smaller pores, stabilizing protrusions and enabling the cell to squeeze through. Similar path choices have been observed in experimental studies with different cell types, where the MTOC is positioned either ahead of or behind the nucleus in pillar parks [21].

## DISCUSSION

Motivated by recent experiments reporting strong correlations between centrosome positioning and migration characteristics and path choices, we presented in this work a mechanistic whole cell model that integrates basic aspects of actin-MT crosstalk, namely growing/shrinking MTs delivering polymerization/contraction signals locally to actomyosin. Our model shows that the position of the centrosome, anterior or posterior to the nucleus, corresponds to different arrangements of the MT array within the cell. This provides different pathways for cell polarization through actin polymerization and myosin contraction signals. Our in-situ results indicate that the position of the centrosome depends on the average lengths of the MTs (Fig. 2**B**). Regular MTs, with an average length of 2/3 of the cell diameter, place the centrosome mostly ahead of the nucleus, whereas long MTs with an average length greater than the cell diameter place the centrosome behind the nucleus in the direction of migration.

The migrating cell can also change from ballistic motion to diffusive motion by adjusting its cortical actin dynamics through changes in the distribution of regulatory signals reaching the cortex. Earlier studies have reported that variations in MT numbers can alter the characteristics of cell migration [48, 49, 53]. Our model also suggests that changes in the number of MTs can lead to different cell migratory behaviors. In a centrosome anterior to the nucleus configuration, an increase in the number of MTs leads to a significant increase in persistence (Fig. 3**D**). In the centrosome posterior to the nucleus configuration, increasing MT numbers initially lead to ballistic migration, but the motion becomes super-diffusive when the number of MTs is too high(Fig. 3**C**).

When cells migrate through complex geometries within tissues, they are faced with obstacles composed of the surrounding extracellular matrix and tissues. These cells have highly branched shapes at junctions where they use protrusions to probe the surrounding narrow channels and choose the most optimal path. Eventually, the entire cell proceeds along one of the protrusions while the others are retracted [54]. MTs play a vital role in the choice of the winning protrusion. The protrusion along which the cell moves is stabilized by the delivery of actin polymerization signals through MTs, whereas the other protrusions are retracted through cortical contraction signals [6].

Experimental studies on *dictyostelium discoideum* and leukocytes moving through pillar parks or Y junction channels suggest that the centrosome position, anterior or posterior to the nucleus, determines the choice of the path of the cell [20, 21]. Further, cells have also been shown to drift away from regions of densely packed obstacles [55]. Our results indicate that when the centrosome is anterior to the nucleus (with regular MT, *l*_*mt*_ = 9.3*µm*), growing MT tips can move into both narrow and wide channels, enhancing actin polymerization activity. The cell’s migration path is determined by which protrusion receives more actin polymerization signals, enabling that protrusion to assist in squeezing the nucleus through the channel. When the nucleus is ahead of the centrosome (with long MTs, *l*_*mt*_ = 16*µm*), it can act as a barrier to the entry of growing MT tips in the narrow channels. The wider channels receive more growing MT tips and actin polymerization signals. This favors the cell moving into the wider channel while the protrusion in the narrow channel is retracted (Fig. 6**G**).

Our simulation results also predict that with the centrosome anterior to the nucleus, cells could migrate more robustly in obstacle parks (Fig. 4**C**). Further, at a Y junction, cells with anterior centrosome position prefer the wider channel only slightly more than the narrow channel. While cells almost always choose the wider channel when the centrosome is posterior to the nucleus (Fig. 6**G**).

Our results can help give insights into how immune cells navigate through the body to reach targets, how cancer cells can start to metastasize, and how cells migrate in synthetic confined geometries. Our model is the first step to integrate basic aspects of actin-MT crosstalk into a mechanistic model for cell migration, and it predicts already a rich variety of emerging self-organized dynamical behavior, including centrosome positioning, switching between random and ballistic movement, and path choice dynamics in confined environments. As such, it contains various simplifications: for instance, our representation of the cell membrane as a closed-loop bead-spring polymer model is limited in capturing the full complexity of the cell membrane. Parameters such as the membrane’s stretching and buckling stiffness could not be directly correlated with the properties of an actual membrane surface. The interaction of dynein motors with MTs at the cortex underlying the cell mem brane also cannot be captured completely through simple spring attachments between discrete membrane and MT beads. Therefore, to align our results with available experimental observations, we chose parameter values that produced the best results.

Our model’s assumption of a consistent average MT length does not fully align with experimental observations. In dendritic cells, where the centrosome is positioned posterior to the nucleus, cell polarization occurs through longer, more stable MTs extending toward the cell front and shorter, less stable MTs directed toward the cell rear [6, 13]. This results in an elliptical morphology, which our model cannot predict, as it lacks a mechanism to stably polarize the cell in a specific migratory mode with an elongated shape and a spatially heterogeneous MT length distribution. Exploring potential mechanisms that lead to such stable polarization and incorporating them into our model would be an interesting direction for future work.

Finally, cells are known to exhibit differential MT stability in response to external chemical or mechanical cues [6, 13, 56–58]. Furthermore, factors such as the arrangement of the extracellular matrix and the density of adhesion molecules can influence the migratory behavior of cells. [59–63]. Cells migrating in 3D extracellular matrix environments often display distinct characteristics compared to those migrating on a 2D glass surface, a distinction that simplistic 2D models fail to capture. Future studies could investigate how cells migrating in both 2D and 3D environments sense external cues, such as chemical signals and the organization of the extra-cellular matrix, to determine their migration direction. Understanding the role of MTs and actin-MT crosstalk in these sensing mechanisms would be an interesting area for further research.

## Supporting information

https://github.com/PinakiNayak13/Modeling_actin_mt_crosstalk_in_migrating_cells

## ACKNOWLEDGMENTS

P.N. was supported by a fellowship from CSIR, India. H.R. acknowledges financial support by the German Research Foundation (DFG), project number 468346334. R.P. thanks IACS for funding and computational facilities.

## APPENDIX

### Short MTs lead to loss of persistence

To test the effect of shortened MTs on cell polarization, cells were simulated with a short average MT length of *l*_*mt*_ = 3*µm*. The cell centroid trajectories revealed that the cell became increasingly prone to turning, leading to a loss of persistence (Fig. 7**A**). The centrosome stayed ahead of the nucleus in the direction of migration (Fig. 7**B**). The mean squared displacement of the cell centroid showed a scaling of 1.1,5, which corresponds to a super-diffusive regime (Fig. 7**C**). Analysis of the velocity direction autocorrelation function indicated that cell migration direction was correlated at small time scales (Fig. 7**D**). With increasing time, the correlation of cell velocity decayed steeply to zero, indicating that the cell was prone to changing direction of propagation with time.

**FIG. 7.**
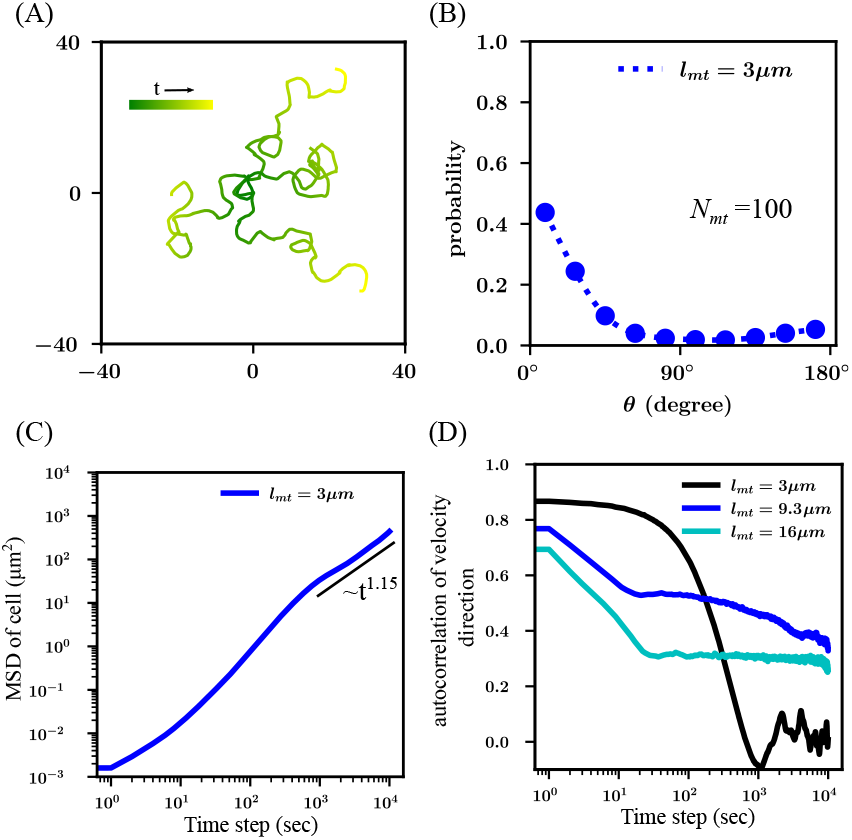
Short MTs frequently cause cells to change migration direction. **(A)** Trajectories of cell centroids with short average MT length of *l*_*mt*_ = 3*µm* at an intermediate membrane and MT stiffness of *k*_*mem*_ = 240*pNµm*^−1^ and *k*_*mt*_ = 400*pNµm*^−1^. **(B)** Probability distribution of angle between direction of cellular motion and nucleus to centrosome vector for *l*_*mt*_ = 3*µm*. **(C)** Mean square displacement of cells showing a *t*^1.15^ dependence corresponding to super diffusive migration. **(D)** Velocity direction autocorrelation function shows velocity direction becomes uncorrelated with time.

### Severely short MTs impair cell migration

The effect of MT depolymerization on the migration was investigated by simulating cells with average MT length severely shortened to *l*_*mt*_ = 1.0*µm*. Our results indicated that the cell failed to establish and maintain any front-back polarization with severely short MTs. Trajectories of cell centroids in Fig. 8**A-B** indicate that the cell centroid did not have a net displacement in the order of the cell diameter throughout the simulation time. We interpret this as an overall failure of the cell migration. Events of migration failure due to severely shortened MTs have been demonstrated in nocodazole-treated cells that depolymerize MTs [64].

**FIG. 8.**
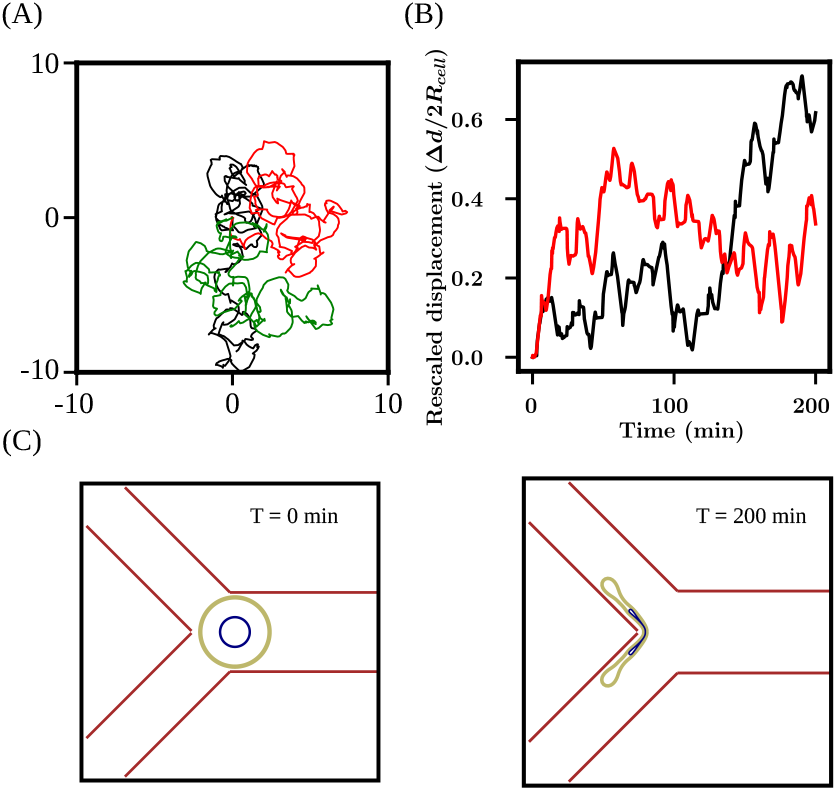
Very short or no MTs lead to migratory failure. **(A)** Typical trajectories of cell centroids with very short MTs (*l*_*mt*_ = 1.0 *µm*) for 0 − 200 *min*. All trajectories start at the origin (0,0). **(B)** Cell centroid displacement from initial position scaled by 2*R*_*cell*_. **(C)** Initial and final configuration of cell without MTs at a Y junction channel. Loss of MTs leads to cell membrane collapse at Y junction.

The functional consequence of complete MT depolymerization for cells migrating in restricted geometries was checked by simulating cells without MTs at a Y junction. A small velocity was given to each cell membrane bead directed inwards into the channel, depending on which channel the bead was in. Cells at a symmetric Y junction (channel width = 10*µm*) failed to migrate into either channel. The cell collapsed at the Y junction in the absence of MTs (Fig. 8**C**).

### Stiffer cell membrane-actin cortex and softer nucleus improves persistence in posterior centrosome configuration

The influence of the cell membrane and underlying actin cortex stiffness on the migration of the cell was investigated by varying the membrane spring constant *k*_*mem*_ in the simulations. For cells with regular MTs (*l*_*mt*_ = 9*µm*) no significant change in migration characteristics were observed. However, cells with long MTs (*l*_*mt*_ = 16*µm*), were found to migrate more persistently in unrestricted geometries with a stiffer cell membrane (Fig. 9**(A)**). Analysis of the local msd exponent *α* revealed that, in the centrosome posterior to nucleus configuration, a stiff cell membrane leads to more directed run phases of the cell (Fig. 9**B,C**). A stiffer cell membrane and actin cortex help in faster retraction of the cell rear through enhanced membrane tension and result in improved directed locomotion of the cell. In obstacle parks, cells with long MTs (*l*_*mt*_ = 16*µm*) were found to show improved migration with a stiffer cell membrane and underlying actin cortex. This was indicated by the presence of higher counts of directed run phases for *k*_*mem*_ = 360*pNµm*^−1^ as compared with *k*_*mem*_ = 120*pNµm*^−1^ (Fig. 9**D,E**). Finally, a softer nucleus was found to improve the capability of the cells to squeeze through narrow pores between obstacles in obstacle parks. For obstacle radius *R*_*obs*_ = 5*µm* and spacing Δ*d* = 20*µm*, cells having long MTs with *k*_*mem*_ = 240*pNµm*^−1^ and *k*_*nuc*_ = 5000*pNµm*^−1^ found it difficult to move through the narrow pores between obstacles resulting in stuck configurations of the cell. However, when the nucleus membrane stiffness was reduced to *k*_*nuc*_ = 2000*pNµm*^−1^ the ability of the cells to migrate through the narrow spacings between obstacles improved, resulting in increased average displacement of the cell centroid with time (Fig. 9**F**).

**FIG. 9.**
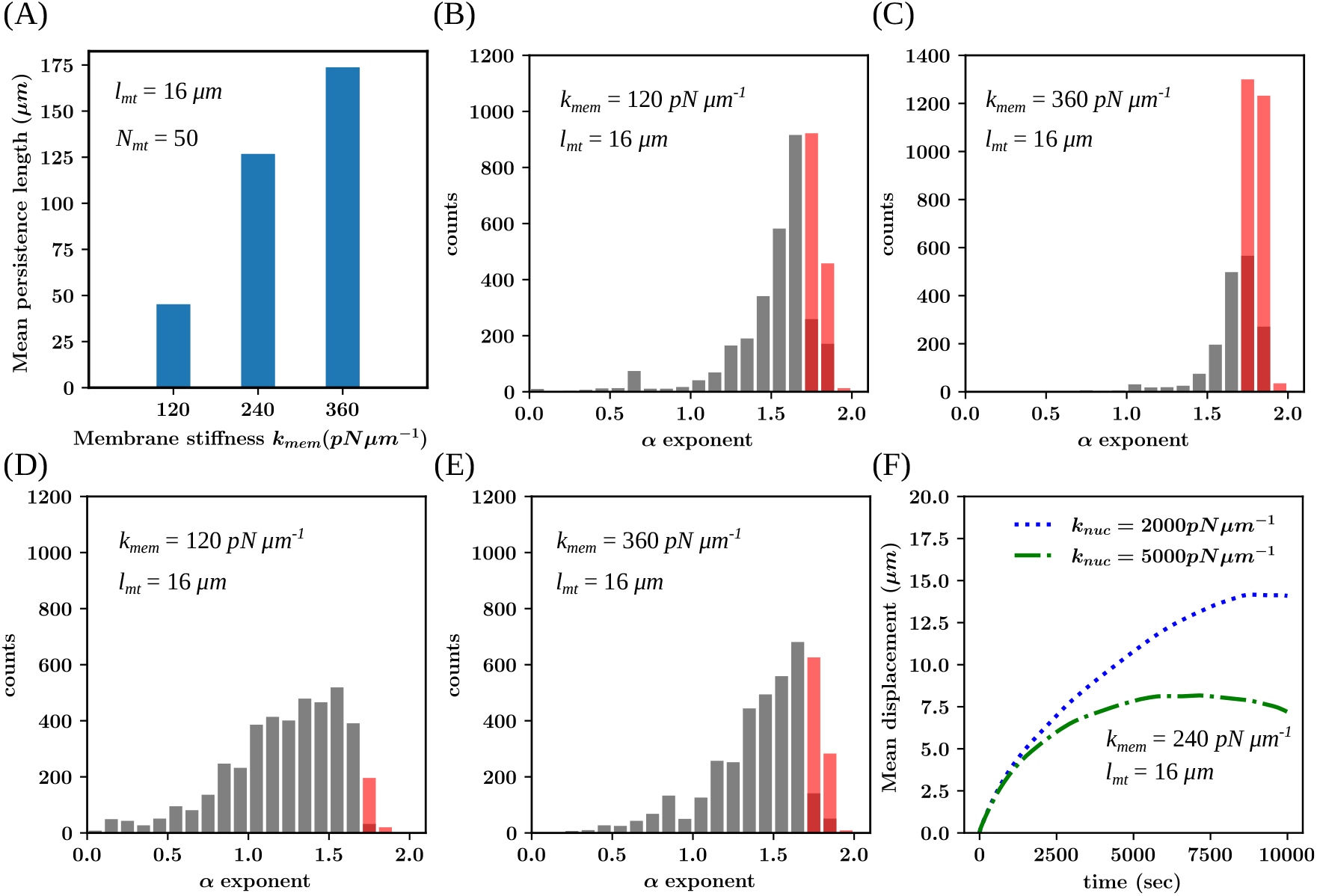
Membrane stiffness affects cell persistence. **(A)** Persistence length of cells with varying membrane stiffness for centrosome posterior to nucleus (*l*_*mt*_ = 16*µm*). **(B)** Distribution of local MSD exponent *α* for freely migrating cells with long MTs and *k*_*mem*_ = 120*pNµm*^−1^. **(C)** Distribution of *α* for freely migrating cells with long MTs and *k*_*mem*_ = 360*pNµm*^−1^. **(D)** Distribution of *α* for migrating cells in obstacle maze with long MTs and *k*_*mem*_ = 120*pNµm*^−1^. **(E)** Distribution of *α* for migrating cells in obstacle maze with long MTs and *k*_*mem*_ = 360*pNµm*^−1^. **(F)** Mean displacement of cells migrating in obstacle maze with long MTs and various nucleus stiffness. All simulations were performed for *N*_*mt*_ = 50.

### Unbiased cells move towards less dense regions in obstacle parks with varying density

Finally, we investigated the behavior of migrating cells within obstacle parks with varying obstacle densities. Cells with regular and long MTs mostly moved towards the region with a sparse distribution of obstacles, with few cells moving toward the region densely packed with obstacles (Fig. 10 **A-C**). This suggested that cells can use their MT-actin crosstalk to navigate towards less restrictive regions. For cells with regular MTs, a greater percentage (≈ 20%) of cells moved towards the denser region as compared with cells with long MTs (≈ 10%). However, cells with regular microtubules were able to move more robustly within the obstacle park, as was also seen for regularly spaced obstacle parks (see Fig. 4).

**FIG. 10.**
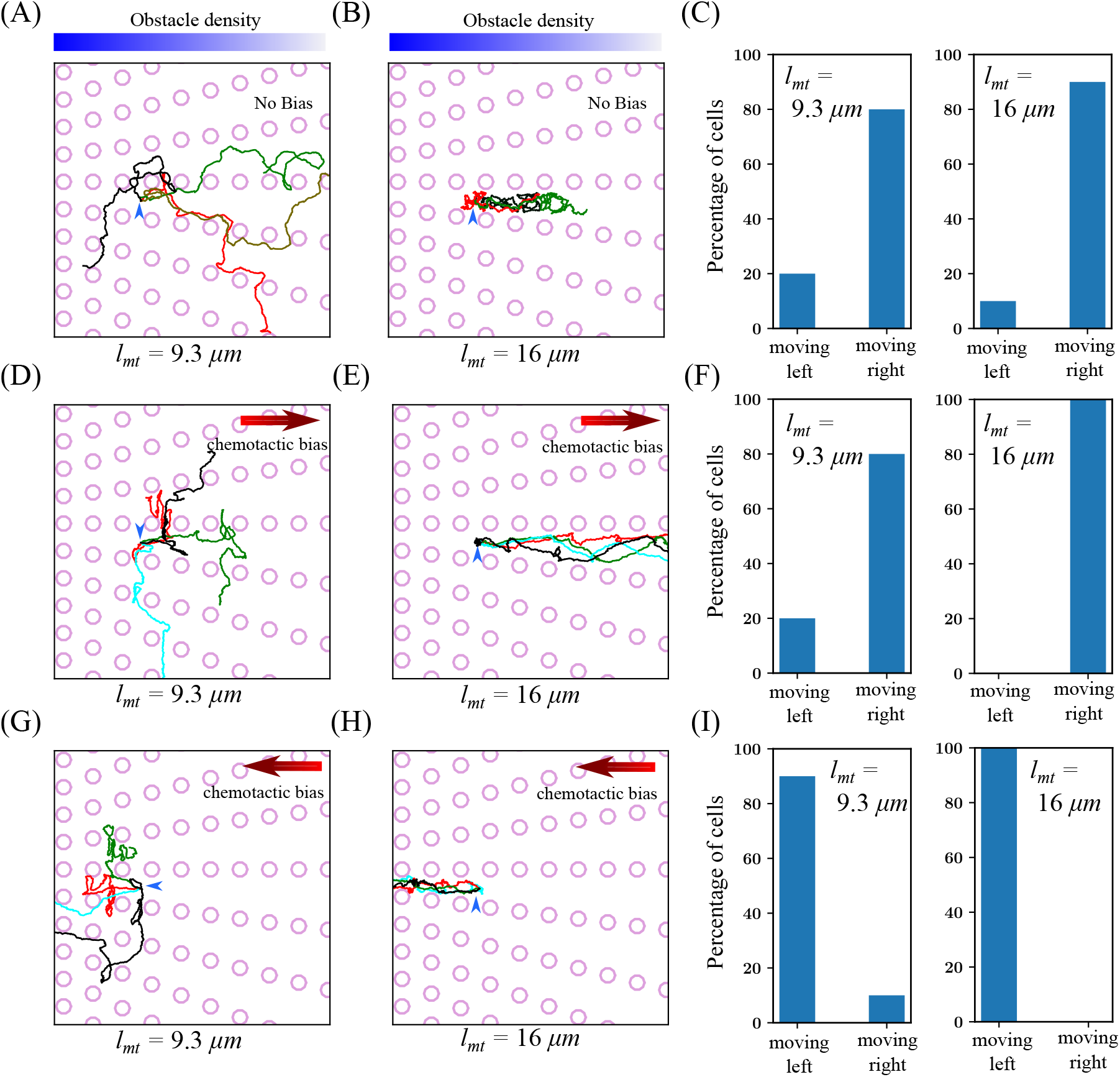
Cell migration in obstacle park with obstacle density gradient. **(A)** Cell trajectories for cells with regular MTs and *N*_*mt*_ = 50, placed in a obstacle park with gradient obstacle density. **(B)** Cell trajectories for cells with long MTs in obstacle park with gradient obstacle density. **(C)** Percentage of cells moving left (towards higher obstacle density region) and cells moving right (lower obstacle density region) for regular and long MTs. **(D)** Cell trajectories for cells with regular MTs and *N*_*mt*_ = 50 in presence of a chemotactic bias towards the right (lower obstacle density region). **(E)** Cell trajectories for cells with long MTs and a chemotactic bias towards the right (lower obstacle density region). **(F)** Percentage of cells moving left and cells moving right for regular and long MTS in presence of chematactic bias towards the right. **(G)** Cell trajectories for cells with regular MTs and a chemotactic bias along the left (higher obstacle density region). **(H)** Cell trajectories for cells with long MTs and a chemotactic bias along the left (higher obstacle density region). **(I)** Percentage of cells moving left and cells moving right for regular and long MTS, with a chemotactic bias towards the left (higher obstacle density region). Blue arrow indicates the initial position of the cell.

Next, we introduced a chemotactic bias on the cell by adding a small velocity to the cell membrane beads in the direction of the bias. The migration of cells with regular MTs was found to be hampered when the chemotactic bias was in the direction of sparsely placked obstacles (Fig. 10 **D**). This indicated that the polarization of the cell due to microtubule-delivered signals failed to align with the chemotactic bias. Therefore the cells demonstrated a reduction in its ability to navigate thorugh the narrow spaces within obstacles. However, most cells moved towards the sparse obstacle density region (Fig. 10 **F**). For long MTs, all the cells moved towards the sparse obstacle density region with the migration becoming more robust (Fig. 10 **E,F**).

The direction of the chemotactic bias was then reversed towards the densely packed region. Cells with regular microtubules mostly moved towards the densely packed region (Fig. 10 **G**). A small percentage (≈ 10%) of cells moved towards the sparsely packed region opposite to the applied chemotactic gradient (Fig. 10 **I**). Cells with long MTs only moved towards the densely packed region, indicating that long MT cells can align themselves better to the applied chemotactic gradient (Fig. 10 **H,I**).

### DETAILS OF THE MODEL

#### Membrane dynamics

The cell and nuclear membrane are modeled as a closed chain of *N*_*mem*_ bead spring units (see Fig. 1**A**) [36, 80, 81]. Each bead is connected to two neighboring beads on either side via a spring, and an angular potential is considered between the angle formed by the three adjacent beads. The nuclear membrane beads also have a Lennard-Jones interaction with the cell membrane beads within a cut-off distance such that the interaction forces are always repulsive.

**TABLE I.** Parameter values used in simulations.

| Symbol | Description | Value | Reference |
| --- | --- | --- | --- |
| $N_m$ | Number of cell membrane beads | 200 | [65] |
| $N_n$ | Number of nuclear membrane beads | 200 | [65] |
| $N_{mt}$ | Number of MTs | 50 - 200 | [66, 67] |
| $N_{dyn}^m$ | Number of dynein motors on cell membrane | 400 | This study |
| $N_{dyn}^n$ | Number of dynein motors on nuclear membrane | 400,600 | This study |
| $R_{cell}$ | Initial radius of cell | $7 \mu m$ | [68, 69] |
| $R_n$ | Initial radius of nucleus | $3 \mu m$ | [70] |
| $v_g$ | MT growth velocity | $0.4 \mu m s^{-1}$ | [71–73] |
| $v_s$ | MT shrink velocity | $0.8 \mu m s^{-1}$ | [72–75] |
| $f_c$ | MT catastrophe frequency | $0.045\text{--}0.06285 s^{-1}$ | [73–75] |
| $f_r$ | MT rescue frequency | $0.04 s^{-1}$ | [73–75] |
| $k_{mem}$ | Cell membrane stiffness | $120\text{--}360 pN \mu m^{-1}$ | [76, 77] |
| $k_\theta^m$ | Cell membrane bending rigidity | $80 pN$ | [76] |
| $l_0^m$ | Rest length of membrane spring attachments | $0.22 \mu m$ | This study |
| $\epsilon_m$ | Strength of LJ interaction between membrane-nucleus | $10.0 pN \mu m^{-1}$ | This study |
| $2^{\frac{1}{6}} \sigma_m$ | Cut-off distance for membrane-nucleus interaction | $0.31 \mu m$ | This study |
| $k_{mt}$ | MT stretching stiffness | $100\text{--}1000 pN \mu m^{-1}$ | [78] |
| $k_\theta^{mt}$ | MT bending rigidity | $0.009 pN$ | [78] |
| $l_0^{mt}$ | Rest length of MT spring attachments | $0.2 \mu m$ | [25] |
| $\epsilon_{mt}$ | Strength of LJ interaction between MT-Cell/Nucleus | $10.0 pN \mu m^{-1}$ | This study |
| $\sigma_{mt}$ | Cut-off distance for MT-Cell/Nucleus LJ interaction | $0.42 \mu m$ | This study |
| $k_d$ | Dynein spring attachment strength | $0.1 pN \mu m^{-1}$ | This study |
| $v_{dyn}$ | Dynein walking speed on MTs | $40 nm s^{-1}$ | [79] |
| $d_{cut}$ | Dynein attachment cut-off distance | $1 \mu m$ | [25] |
| $k_c$ | Nucleus to cell membrane linear coupling strength | $0.25 pN \mu m^{-1}$ | This study |
| $l_p S_0 r_0$ | Actin polymerization velocity in absence of polymerization signals | $0.1 \mu m s^{-1}$ | This study |
| $C_p$ | Actin polymerization velocity increment per growing MT tip | $0.1 \mu m s^{-1}$ | This study |
| $F_0$ | Cortical acto-myosin contraction force in absence of contraction signals | $1.25 - 2.5 pN$ | This study |
| $C_d$ | Cortical contraction force increment per shrinking MT tip | $2.5 pN$ | This study |
| $\zeta_c$ | Effective coefficient of viscosity of cell membrane beads | $20 pN \mu m^{-1} sec$ | This study |
| $\zeta_n$ | Effective coefficient of viscosity of nucleus membrane beads | $25 pN \mu m^{-1} sec$ | This study |
| $\zeta_{mt}$ | Effective coefficient of viscosity of MT beads | $5 pN \mu m^{-1} sec$ | [25] |
| $\zeta_{mt}$ | Effective coefficient of viscosity of centrosome | $10 pN \mu m^{-1} sec$ | This study |

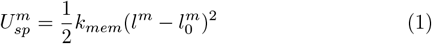

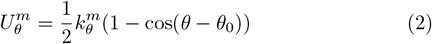

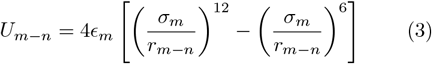

Here, *l*^*m*^ denotes the stretched length of the spring joining two adjacent membrane beads, 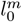 denotes the rest length of the spring joining two adjacent membrane beads, and *r*_*m*−*n*_ denotes the distance between a membrane bead and nuclear membrane bead. The equation of motion for a cell or nuclear membrane bead then follows as,

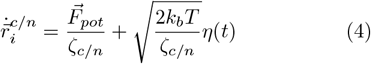

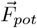 represents the sum total of all forces on the beads due to the potentials considered in equations (1-3). *ζ*_*c/n*_ is the coefficient of viscosity of the membrane/nucleus beads and *η*(*t*) is a Gaussian noise with zero mean and ⟨*η*(*t*)*η*(*t*^*′*^)⟩ = *δ*(*t* − *t*^*′*^).

#### MT dynamics

MTs are modeled as open-ended bead spring units with their minus ends anchored at the centrosome [78]. The MT beads are connected to adjacent beads on either side via a spring and an angular potential is considered between the angle formed between three adjacent beads in an MT. The dynamic instability of the MT is incorporated into the model by adding (or removing) beads at the plus end of a growing (or shrinking) MT at intervals of *t*_*mt*_ timesteps (see Fig. 1**B**). New beads are added to growing MTs in a given time step only if the new bead falls within the cell and outside the nucleus. The dynamic instability of MTs is governed by four parameters, namely, growth velocity (*v*_*g*_), shrink velocity (*v*_*s*_), catastrophe frequency (*f*_*c*_), and rescue frequency (*f*_*r*_) [74, 75, 82]. An increase in catastrophe frequency (*f*_*c*_), or a decrease in rescue frequency (*f*_*r*_) reduces the average MT length. Similarly, an increase in rescue frequency (*f*_*r*_), or a decrease in catastrophe frequency (*f*_*c*_) increases the average MT length in the cell.

The membrane beads and MT beads have Lennard-Jones interaction between them within a cut-off distance such that the interaction forces are always repulsive.

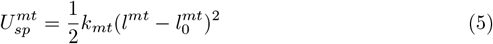

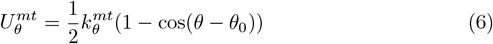

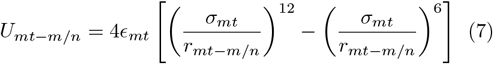

Here, *l*^*mt*^ denotes the stretched length of the spring joining two adjacent MT beads, 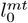 denotes the rest length of the spring joining two adjacent MT beads, *r*_*mt*−*m/n*_ denotes the distance between a MT bead and a membrane bead (or a nuclear membrane bead). The equation of motion for an MT bead reads,

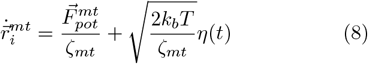

#### Dynein dynamics

Dynein motors are placed randomly on membrane beads (or nuclear membrane beads) and scanned for MT beads near them. Upon encountering MT beads within a cut-off distance *d*_*cut*_, dynein motors form spring-like bonds with them. After each time step, the dynein motors can shift to the adjoining MT bead toward the negative end of the MT if the adjoining bead is within the cutoff radius *d*_*cut*_. The dynein walking speed on the MTs is given by *v*_*d*_*yn*. Dynein pulling forces are modeled as simple spring forces between the membrane (or nuclear membrane) beads and MT beads.

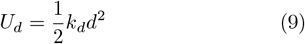

Where *d* is the distance between the centers of the membrane (or nuclear membrane) bead and the MT bead.

#### Actin dynamics

We modeled the effects of actin network dynamics as protrusive and contractile forces on the membrane beads. The protrusive forces caused by actin polymerization are considered to add a component to the velocity of the membrane beads directed in the outward normal direction [33]. This outward component of the velocity depends on the average local orientation *S*_*act*_ of the actin filaments in the vicinity of the membrane bead and the local actin polymerization rate *r*_*p*_. The net outward velocity added to the membrane beads is then *l*_*a*_*S*_*act*_*r*_*p*_, where *l*_*a*_ is the diameter of individual actin filaments. We consider two types of contractile forces on the cell membrane beads due to actomyosin activity. Myosin activity on counter-oriented long actin filaments inside a network in the crossover region between the protrusion and cell body generates contractile forces on the protrusion. The crossover region consists of actin filaments associated with the nucleus along with actin filaments associated with the adhesions in the protrusion. The strength of this contractile force depends on the actin retrograde flow velocity 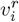 at the protruding membrane bead. To evaluate the net velocity change of the protruding membrane beads due to cortical actin activity, we follow [33], and equate the net change in bead velocity to the difference between the outward actin polymerization velocity and retrograde flow velocity.

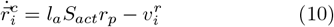

The retrograde flow velocity can be assumed to be driven by the membrane tension *f*_*τ*_ and the contraction force *f*_*c*_ at the membrane due to coupling with the nucleus through actin filaments and myosin motors. It can be shown that *f*_*c*_ and *f*_*τ*_ vary linearly with the length of the protrusion [33, 42, 45, 46].

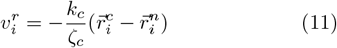

Where *k*_*c*_ represents the strength of the linear coupling between the membrane beads and nuclear membrane beads. The protrusion length is the distance 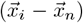 between the membrane bead and the nuclear membrane bead and *ζ* is the effective friction coefficient on the membrane beads due to focal adhesions. We also consider a local contractile force *F*_*c*_ due to myosin activity in the cell cortex along the cell membrane that does not couple with the nucleus, directed opposite to the outward normal at the membrane. The final equation of motion for the membrane beads then reads as follows:

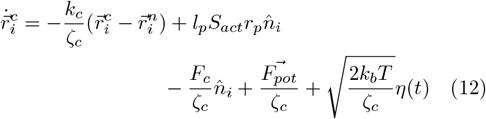

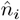 is a unit vector pointing along the outward normal along the membrane bead. *F*_*pot*_ is the sum of all the forces due to the interaction potentials between the membrane and the nucleus and MT beads. We assume that the values of *S*_*act*_ and *r*_*p*_ depend on actin polymerization cues that are supplied to the region near the membrane beads by polymerizing MTs and their values are calculated according to,

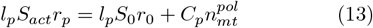

Where *l*_*p*_*S*_0_*r*_0_ is the component of the outward polarization velocity in the absence of polarization cues delivered by the growing MT tip, 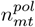 is the number of polymerizing MTs in the vicinity of the membrane bead and *C*_*p*_ is the component of protrusion velocity added per polymerizing MT. The strength of the contraction force *F*_*c*_, due to myosin activity at the cortex, is considered to depend on contraction cues provided by depolymerizing MT tips near the membrane beads and varies according to :

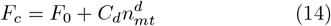

Where 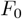 is the strength of the contraction force in the absence of any MT-supplied contraction cue. *C*_*d*_ is the net contraction force added per depolymerizing MT present near the membrane bead, and 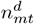 is the number of depolymerizing MTs near the membrane bead. Finally, the equation of motion of the nucleus beads reads,

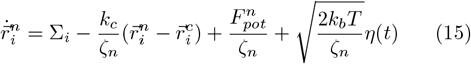

Where 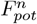 represents the sum of all the forces on the nucleus bead due to the steric and dynein interactions considered in our model.

### SIMULATION METHOD

All simulations were initialized with a circular cell and nucleus and the centrosome was placed randomly within the cell. All MTs were initialized to be growing and their directions of growth were chosen randomly. The simulation time step was taken to be 0.1*sec*. The beads were added or removed from the MT tips after each *N*_*pol*_ time step according to their growth or shrinkage state. The probabilities for MT catastrophe or rescue were calculated after every *N*_*pol*_ time steps and the MT state was updated accordingly. The number of growing or shrinking MT tips near a membrane bead was checked at every time step and actin polymerization or cortical contraction signal strengths were calculated. Dynein motors were placed on the cell membrane and nuclear membrane beads randomly, and positions of attachments were searched on nearby MTs at every time step. Upon finding possible sites of attachments, dynein motor bonds were established between the membrane and MT beads with a certain probability. Established dynein bonds were checked for turnover and bonds were detached according to the turnover probability at every time step. For the force dynamics, we considered a relaxation timestep of 0.001*sec*. Forces on every component in the model were calculated at every relaxation time step and their positions were updated accordingly.

The codes were developed in C using OpenMP. The data analysis and plots were done in Python and Gnuplot. The computational time for a single simulation running on 20 processors (Intel Xeon CPU, having a clock speed of 2 GHz and RAM 64 GB) was in the range of 10-20 hours.

### DATA ANALYSIS

#### Persistence length

The local directional persistence, or the ability of the cell to maintain its direction of motion, is quantified as *p* = *cosθ* with *θ* being the angle between the instantaneous velocity directions at two time steps. The persistence length *l*_*p*_ can then be calculated from 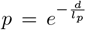,where *d* is the displacement of the cell between the two time steps. The local persistence length *l*_*p*_ was evaluated at consecutive time steps throughout the motion of the cell and its average was calculated as the mean persistence length ⟨*l*_*p*_⟩ [52].

#### Local mean square displacement and velocity standard deviation analysis of cell trajectories

Local mean square displacements were evaluated by considering a rolling time window of *N*_*t*_ = 30 points at consecutive time steps 300 seconds apart. At every time step *t*_*i*_, the local mean square displacement 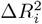 was evaluated as a function of the time lag *τ*_*m*_ = *mδt* as [51],

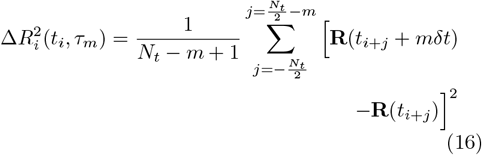

where *δt* is the time between two rolling window points and **R**_*i*_ = (*X*_*i*_, *Y*_*i*_) are the coordinates of the cell centroid. The total duration of the rolling window is *T* = *N*_*t*_*δt*.

The standard deviation of the velocity was calculated at consecutive time steps from the values of the velocity direction *ϕ*_*i*_(*t*_*i*_) as [51],

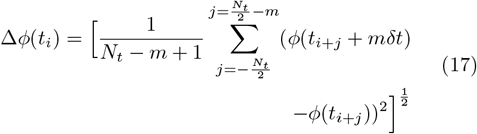

the value of *m* was chosen such that *mδt* = *T/*4.

The local mean square displacement 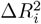 was assumed to scale with the time lag *τ*_*m*_ as,

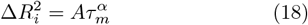

The value of *α* = 1 the motion of the cell is completely diffusive and for a value of *α* = 2 the motion of the cell is ballistic. We considered the motion of the cell in two separate phases. For *α >* 1.7 and Δ*ϕ <* 0.9, the cell migration was considered to be in the directed motion phase, otherwise, the cell motion was considered to be in the random migration phase.

