## Supplementary material for "Modelling Actin-Microtubule Crosstalk in Migrating Cells": https://github.com/PinakiNayak13/Modeling_actin_mt_crosstalk_in_migrating_cells

Movies depicting cell migration in free and restricted environments.


(Movie. 1) Persistent cell migration for N_mt  = 100 and l_mt  = 9.3 μm ; Persistent cell migration for N_mt = 50 and l_mt = 16 μm.


(Movie. 2) Diffusive cell migration for N_mt = 100 and l_mt = 3 μm ; Diffusive cell migration for N_mt = 200 and l_mt = 16 μm.


(Movie. 3) Cell migration in an obstacle park with obstacle spacing d_obs  = 20 μm and obstacle radius R_obs = 4 μm.


(Movie. 4) Cell migration in an obstacle park with obstacle spacing d_obs  = 20 μm and obstacle radius R_obs = 5 μm.


(Movie. 5) Cell migration in path choice device with 4 channels of width 2 μm, 4 μm, 6 μm, 8 μm.
